# A high throughput assay to detect enzymatic polyethylene oxidation

**DOI:** 10.1101/2025.07.23.666384

**Authors:** Ross R. Klauer, Mekhi Williams, Darien K. Nguyen, Megan Tarr, Dionisios G. Vlachos, Kevin V. Solomon, Mark A. Blenner

## Abstract

Biological plastics deconstruction and upcycling have emerged as a sustainable alternative to traditional recycling technologies for plastics waste. The discovery and engineering of efficient thermostable poly(ethylene terephthalate) (PET) hydrolases has made biological PET recycling possible at scale; however, enzymes for non-PET plastics, which account for approximately 70% of all plastics produced, remain largely undiscovered. To accelerate the discovery of such enzymes, a high throughput screening (HTS) platform is needed. Here, we develop a HTS liquid-based assay to detect one of the first committed steps of polyolefin degradation, oxidation of the C-H bond to an aldehyde. We test 4-hydrazino-7-nitro-2,1,3-benxoxadiozole hydrazine (NBD-H), which reacts with generated aldehydes to form a fluorescent hydrazone, on oxidized low-density polyethylene (LDPE) films. Hydrazone generation correlated well with established carbonyl index metrics for polymer oxidation (R^2^ = 0.97). Moreover, we demonstrate that the probe reliably identifies LDPE-active dye decolorizing peroxidases (DyPs) that generate aldehydes on LDPE films, serving as effective screen as demonstrated by a receiver operating characteristic area under the curve of 0.95. Due to the rapid fluorescent readout and parallelization in microarray plates, this assay enables screening thousands of enzymes in 24 hours compared to time-consuming established approaches, accelerating discovery of enzymes that catalyze the first step of polyolefin biodeconstruction.

## 1. Introduction

Enzymes for deconstruction and upcycling of polyolefins, which account for 70% of globally produced plastics [1], remain largely undiscovered [2]. Dye decolorizing peroxidases (DyPs) [3], phenol oxidases [4], and multi-copper oxidases[5] have been reported to oxidize and degrade polyethylene (PE), the most abundant polyolefin. Other enzymes such as laccases[6], manganese peroxidases[7], alkane hydroxylases[8], and Baeyer-Villiger monooxygenases[9] have been linked to PE deconstruction but their known activity is limited to chemically pre-oxidized or very low molecular weight substrates. This paucity of characterized polyolefin active enzymes is largely due to a reliance on time-consuming indirect materials characterization methodologies for validation of enzyme activity, namely Fourier transform infrared spectroscopy (FTIR), Raman spectroscopy, and gel permeation chromatography (GPC)[10]. In the absence of rigorous controls, these methodologies can lead to false positive or irreproducible claims of enzymatic polyolefin deconstruction due to misinterpretation of the complex spectroscopic and molecular weight distribution signals generated[11]. Thus, the field has called for the use of unambiguous isotope labeled plastics, more rigorous controls and/or more careful analysis of spectroscopic or GPC data to confirm enzymatic plastics deconstruction[10]. Nonetheless, these approaches are expensive, time consuming and too low throughout to screen large numbers of enzymes.

Therefore, there is a need for a highly parallelizable and automatable screening methodology to accelerate polyolefin deconstructing enzyme discovery. Such high throughput screening methods greatly accelerated the discovery and development of poly(ethylene terephthalate) (PET) hydrolases, leading to enzymes that have been developed for industrial scale PET deconstruction for recycling or upcycling[12,13]. PET hydrolases produce mono(2-hydroxyethyl) terephthalate (MHET) and bis(2-hydroxyethyl) terephthalate (BHET) that are detectable via high-performance liquid chromatography (HPLC) [13–15], allowing for high throughput testing of many candidate PET hydrolases in parallel. Conversely, biological deconstruction of polyolefins results in a complex mixture of deconstruction products[16] that are poorly defined and, thus, challenging to separate, detect and quantify via chromatographic approaches. Polyolefins are hypothesized to follow canonical alkane catabolism, where alkane chains are first oxidized to form hydroxyls that are then oxidized into carbonyl groups[3,16–18]. These oxidation events can happen anywhere on the polyolefin hydrocarbon backbone, yielding highly variable deconstruction products[16,17]. Moreover, these oxidation sites are influenced by polymer crystallinity, molecular weight, and additive packages that dictate enzymatic access to polymer chains [19–21]. That is, plastics with the same backbone structure, or resin code, need not generate the same deconstruction products. Due to the multi-step deconstruction process and deconstruction product variability, high throughput screening methodologies should be developed based on chemical changes to the polymer backbone in the deconstruction process, which are agnostic to specific deconstruction product generation.

In this work, we develop a fluorescence-based assay for oxidative enzymes that initiate deconstruction of low-density polyethylene (LDPE). We use a highly selective 4-hydrazino-7-nitro-2,1,3-benxoxadiozole hydrazine (NBD-H) chemistry to detect aldehydes on LDPE films. NBD-H is used for the high throughput detection of aldehydes on hydrocarbon substrates as it reacts with them to generate fluorescent hydrazones for easily quantifiable detection[22]. The use of NBD-H on insoluble LDPE was validated using cold-plasma oxidized LDPE films as a positive control[23]. Moreover, NBD-H signal correlated well with carbonyl index measured via FTIR, demonstrating its use as a semi-quantitative metric for LDPE oxidation detection. We further demonstrate that NBD-H can be used to identify enzymatic activity by detecting LDPE oxidation by previously reported LDPE-active DyPs[3]. Our work establishes a novel platform for screening LDPE oxidation activity that can be readily parallelized and automated for accelerated discovery of enzymes involved in polyolefin deconstruction.

## 2. Materials and Methods

### 2.1 NBD-H reaction with 3-pentanone and pentanal

A 300μL 29:1 Acetonitrile-aqueous reaction volume consisting of 33.3 3mM aqueous HCl, 4.5 or 0.45 ppm pentanal or 3-pentanone, and 0.033 μM NBD-H were added to each well in a black bottom, black wall 96-well plate. The reaction mixture was incubated at 40°C for 30 min before reading the fluorescence at excitation 470 nm and emission 530 nm using a BioTEK synergy neo2 multi-mode platereader and gen5 software (version 3.11). Multiple concentrations of NBD-H were used in the concentration optimization experiment, but all other reactions used 0.033 μM NBD-H.

### 2.2 Plasma treatment of LDPE films

Cold, atmospheric plasma was generated in a parallel-plate dielectric barrier discharge (DBD) reactor in an airtight chamber (10 in × 10 in × 10 in, Sanatron). Two stainless-steel electrodes attached with quartz plates (Ø 75 mm × 2 mm), acting as the dielectrics, were positioned parallel to each other and separated using Teflon spacers (Ø 6 mm × 2 mm). The top electrode configuration (Ø 65 mm × 15 mm) was connected to a sinusoidal AC power supply (PVM500), while the bottom electrode configuration (Ø 50 mm × 15 mm) was grounded. During plasma treatment, LDPE films were placed on the quartz plate attached to the ground electrode, and the DBD chamber was purged with 1% O_2_ in He gas for 20 min to remove the air. He/O_2_ plasma was generated at a discharge frequency of 22.5 kHz and a peak-to-peak applied voltage of 4.5 kV, achieving a ∼2.2 W dissipated power.

### 2.3 FTIR

In-house stripped LDPE films were analyzed for changes to the inherent functional groups using FTIR (Thermo Scientific Nicolet iS5 FTIR Spectrometer, Pittsburgh, PA). Spectra for each powdered sample were recorded on a diamond crystal, attenuated total reflectance (ATR) cell. Spectra were recorded in the range of wavelengths 4000–500 cm^−1^ with a minimum of 32 scans and a spectral resolution of 0.482 cm^−1^. Films were rinsed with 500 μL of water followed by 500 μL of 70% ethanol by vortexing for 1 minute at 200 rpm. If FTIR spectra indicated microbial or protein contamination, 70% ethanol on a cotton swab was lightly wiped over the surface to clean residual contamination. Carbonyl indices were calculated as previously reported[3].

### 2.4 Aldehyde peak deconvolution

OriginLab (version 10.15) Multiple Peak Fit tool was used to deconvolute aldehyde and ketone peak areas in FTIR spectra for Figure 6B correlation calculations. The aldehyde peak was set to a center of 1737 cm^-1^ and ketone peak area was set to a center of 1715 cm^-1^ [33]. Baseline was set by subtracting a straight line in OriginLab between 1700 1800 cm^-1^ and a Gaussian fit was used to fit both the ketone and aldehyde peaks. Area under the aldehyde peak was used in Fig. 6B analysis

### 2.5 XPS

XPS was performed using a Thermo Fisher K-Alpha instrument equipped with an Al (K) X-ray. Spectra were obtained with pass energies of 20, 10, and 20 eV for the survey, C1s, and O1s spectra, respectively, at 100 ms dwell time. XPS spectral scans, C1s, and O1s scans were collected on four locations on each film, using the following settings: C1s spectra were analyzed assuming a C-C, C-H peak at 285.0 eV and then peak shifts of 1.5 eV, 3.0 eV, and 4.5 eV for C-O, C=O, and O-C=O peaks, respectively [34]. All spectral analysis was assessed by averaging the results from survey scans on four randomly selected points each film.

### 2.6 NBD-H assay on LDPE films

Each LDPE film (Sigma Aldrich Chemical Company, St. Louis, MO, USA; catalog number 428043 – 250 g, stripped of additives and pressed per previously reported methods13 was cut into a circular shape to fit flat at the bottom of a black 96-well plate. The base of a 200 μL pipette tip (VWR. Radnor, PA, Catalog number 76322-150) served as a stencil for cutting the films (or a 5 mm × 5 mm square in Fig. 3D). Each film was dosed with 0.1mM hydrogen peroxide and 10 μL of 1g/L enzyme in 50 mM phosphate buffer, pH4 in 30 μL reaction doses as previously reported13. The films were then allowed to incubate at 30°C overnight to allow the films were dry. Additional doses were performed as needed for each experiment. After the final dose dried, each well was filled with 300μL of a 29:1 Acetonitrile-aqueous reaction mixture consisting of 33.33mM aqueous HCl and 0.033 μM NBD-H. The plate was incubated at 40°C for 45 min (films were removed after 0, 10, 20, 30, 40, and 60 min for optimization of kinetics in Fig. 3B) with a cover on the plate to reduce the evaporation of acetonitrile. The reaction mixture was then removed, and the wells were washed by aspirating 300 μL of acetonitrile. This wash step was repeated twice before adding 300μL of acetonitrile to the wells and reading fluorescence on the plate reader at excitation 470 nm emission 530 nm.

### 2.7 Best fit analysis for sigmoid function on Figure 3B

A sigmoid function, Equation 1, was used to develop the best fit curve for the data.

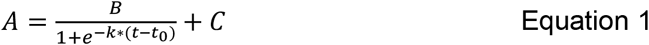

t values were the time values in data in Figure 3B and A values were the fluorescence values from Figure 3B. Excel’s solver function was used to minimize the difference of the true A values vs. the sigmoid function A values by varying B, k, and C. With initial guesses of b = 50, k = 0.1, and t_0_ = 0, the solver provided values of B = 41.04, k = 0.21, t_0_ = 18.86.

### 2.8 DyP expression

For protein expression, *E. coli* BL21 strains containing the DyP on a pET-28a vector[3] were inoculated at 37ºC until they reached an optical density between 0.5 – 0.9 OD600, at which point 0.1 mM IPTG was added to each culture for induction of protein expression. Cultures were then grown at 18 °C for 18-22 hours overnight to express protein, which were then visualized via SDS-PAGE to confirm expression. Sequences for all enzymes are reported in previously published work[3].

### 2.9 Protein purification

After expression, the cultures were pelleted and frozen to prepare for protein extraction using Solulyse bacterial protein extraction reagent (Genlantis Inc. San Diego, CA, USA. Catalog number L100500) per the manufacturer’s instructions. DyPs were purified using His-tag purification with Ni-NTA magnetic beads (New England Biolabs, Ipswich, MA, USA Catalog number S1423L). The procedure provided by NEB per the manufacturer instruction for Ni-NTA magnetic beads was used except for the use of 100μL of magnetic beads for expression cultures of 50 mL or greater. Additionally, 200μL of elution buffer was used instead of 100 μL when expression cultures were 50 mL or greater. Protein concentrations were determined using a Bradford assay and were normalized to 1 g/L by diluting with pH 4 50 mM phosphate buffer.

### 2.10 DyP testing on LDPE films

For activity screening, proteins were heterologously expressed in *E. coli* BL21(DE3) and purified via Ni-NTA magnetic bead purification. Enzymes were dosed onto LDPE films in two doses and allowed to dry overnight between each dose prior to screening for oxidation via NBD-H or FTIR. To screen enzymes for activity, 10 μL of 1.0 g/L enzyme was dosed onto a film an LDPE film stripped of additives from (Sigma Aldrich Chemical Company, St. Louis, MO, USA; catalog number 428043 – 250 g) per previously reported methods or additive-free LDPE films (0.035 mm thick, biaxially oriented LDPE films, purchased from Goodfellow Cambridge Limited, Huntingdon, England. Catalog number LS580568) [3]. Enzymatic reactions of DyPs were carried out at 1 mM H_2_O_2_ (Sigma Aldrich Chemical Company. St. Louis, MO, USA Catalog number H1009 – 500ML) and a pH of 4.0 as a result of enzyme optimization studies [35–37]. Inactive controls were denatured using 1 μL of beta-mercaptoethanol for every 10 μL of 1 mg/mL enzyme. Two doses of this nature were performed, with the last dose allowed to react and dry overnight for 16 hours. Enzyme re-dosing was deemed necessary as a single dose saturating the surface proved insufficient for deconstruction. Moreover, the film surface dried as liquid evaporates, requiring enzyme to be re-dosed to sustain the reaction. Plastic films were then washed in water and ethanol, air dried, and analyzed via FTIR or NBD-H to monitor chemical changes. Unless the name was explicitly stated otherwise, ‘DyP’ tested (Figure 5) was Cp+CNHL DyP.

### 2.11 Pyrogallol assay to confirm DyP inactivity

Pyrogallol (Sigma Aldrich Chemical Company. St. Louis, MO, USA Catalog number P0381 – 25 g) was used as a standard substrate for measuring enzyme activity of peroxidases. The enzyme assay used followed the ‘Enzymatic Assay of Peroxidase (EC 1.11.1.7)’ protocol from Millipore sigma (86). Briefly, 0.027% v/v hydrogen peroxide was added with 0.5% w/v pyrogallol and 0.75 units of peroxidase for the reaction. The generation of purpurgallin was measured using absorbance at 420 nm. Inactive controls were denatured using 1 μL of beta-mercaptoethanol for every 10 μL of 1.0 g/L enzyme.

### 2.12 Statistical testing

All statistical tests were performed using a one-tailed, heteroscedastic t-test between groups of independently prepared biological sample replicates.

## 3. Results

### 3.1 NBD-hydrazine can be used to detect aldehydes selectively

Polyolefins require oxidative chemistries to functionalize highly stable C-C bonds and activate them for biodeconstruction[16]. Identification of these oxidation events is crucial for identifying microorganisms and enzymes that oxidize and deconstruct polyolefins. We thus evaluated NBD-H as a candidate probe for carbonyl detection on oxidized LDPE films, which was selected as the plastic of interest due to its 1) prevalence[1], 2) simple carbon-carbon backbone that is the foundational structure for all polyolefins [1,16], and 3) low crystallinity relative to HDPE that makes it more bioavailable [19,20]. NBD-H was selected due to its extremely high selectivity for aldehydes[24] that we hypothesized would limit non-specific reactions with carbonyl groups common in biomolecules present in the sample that could lead to false positive signals. Moreover, enzymatic LDPE oxidation by DyPs leads to approximately a 50/50 mixture of aldehydes and ketones, making aldehydes a good proxy for LDPE oxidation activity[3]. NBD-H produces a quantifiable signal by selectively reacting with aldehydes to form fluorescent hydrazone derivatives (Fig. 1A). We confirmed NBD-H selectivity for aldehydes by demonstrating a fluorescent response in the presence of pentanal and little to no signal in the presence of 3-pentanone (Fig. 1B). This hydrazine signal was dose-dependent, allowing for quantification of pentanal concentration(Fig. 1B), in agreement with prior studies with aliphatic aldehydes[22].

**Figure 1.**
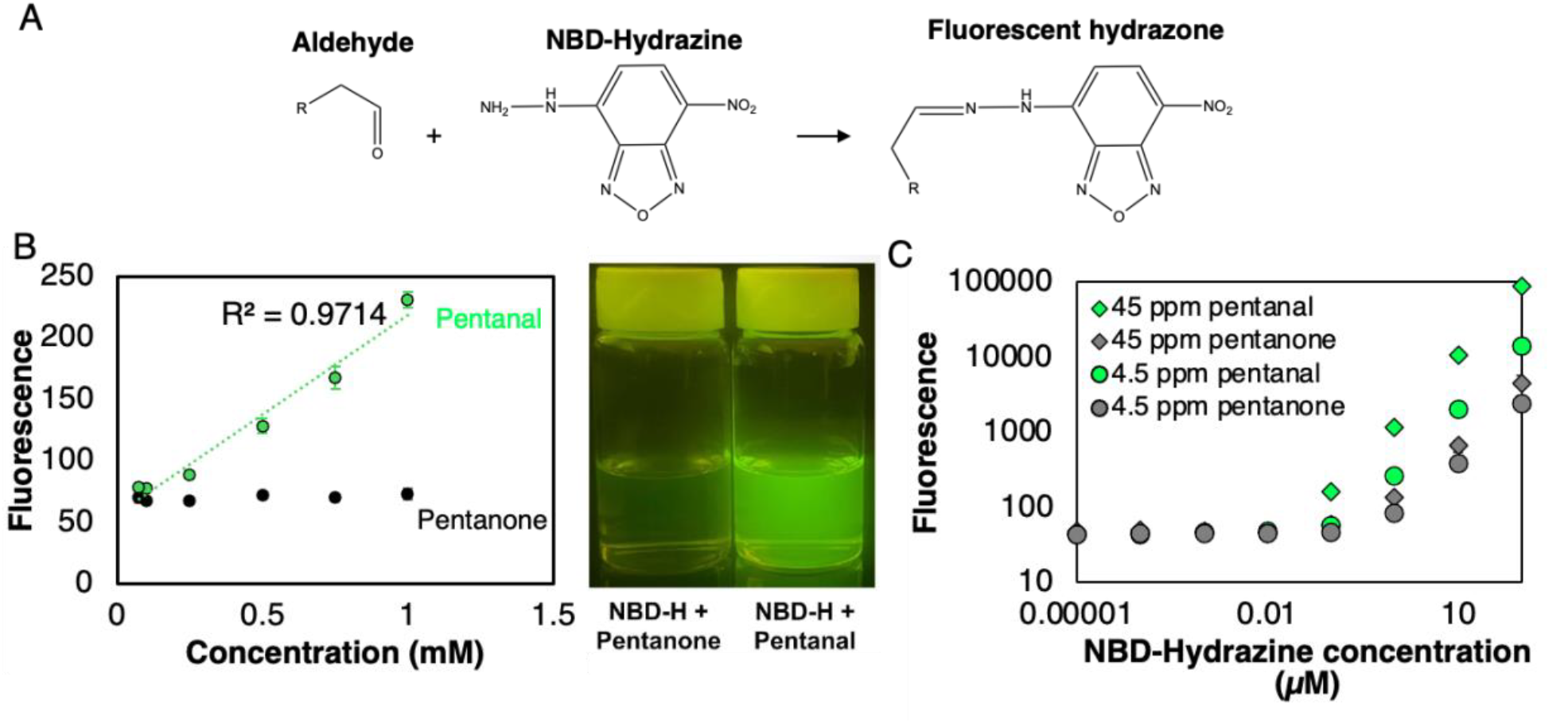
NBD-H selectively detects aldehydes. (A) Chemical reaction between NBD-H and the aldehyde produces a fluorescent hydrazone derivative. (B) NBD-H signal is specific to aldehydes over ketones. NBD-H fluorescence at excitation 470 nm and emission 530 nm on NBD-H after reaction with pentanal and 3-pentanone and 0.1 mM NBD-H (left). Example of NBD-H signal reacted with 1 mM 3-pentanone and 1 mM pentanal (right). Error bars represent standard error of independent triplicate measurements and R^2^ value was calculated via linear regression. (C) NBD-H concentration optimization of 4.5 ppm and 45 ppm pentanal (green) and negative control 4.5 ppm and 45 ppm 3-pentanone (grey). Error bars represent standard error of independent triplicate measurements

To increase assay sensitivity, we optimized the concentration of fluorogenic NBD-H. We found that NBD-H produces an auto-fluorescent signal at high concentrations (> 0.1 μM) in acetonitrile (Fig. 1C). Thus, we decreased NBD-H loading concentrations to determine where pentanal can be detected with limited background interference, assessed using equivalent concentrations of 3-pentanone as a negative control. NBD-H concentrations up to 0.01 μM were too dilute to detect a signal change in the presence of pentanal (Fig. 1C). Concentrations of at least 1 μM led to a false positive ketone signal, thereby increasing assay noise (Fig. 1C, Supplementary Fiig. 1). 0.1 μM NBD-H loading led to the maximum signal:noise ratio at pentanal concentrations of 4.5 and 45 ppm (Fig. 1C, Supplementary Fiig. 1). Therefore, 0.1 μM NBD-H was used in all assays moving forward.

### 3.2 NBD-hydrazine detects carbonyls on chemically oxidized LDPE films

We generated positive control LDPE samples for NBD-H oxidation detection by oxidizing LDPE films using cold plasma. LDPE films were stripped of additives as reported previously[3] to ensure any oxidation signal was a result of cold plasma treatment and not from plastic additives. Cold plasma treatment was selected because oxygen radicals generated in plasma can react with LDPE films, introducing oxygen functional groups on the surface[25]. This enables rapid oxidation of the surface of LDPE films (ranging from nanometers to microns) while preserving the molecular weight distribution of the bulk polymer[26,27], emulating biochemistries performed by LDPE- oxidizing enzymes. Cold, atmospheric plasma was generated in a parallel-plate dielectric barrier discharge (DBD) reactor in an airtight chamber where LDPE films were treated with He/O_2_ plasma generated at a discharge frequency of 22.5 kHz and a peak-to-peak applied voltage of 4.5 kV, achieving a ∼2.2 W dissipated power (Fig. 2A). LDPE films were oxidized by plasma treatment, where both global oxygen content and carbonyl content increased as a function of treatment time, producing 4.4%, 5.4%, and 5.9% C=O bonds as a result of 5-, 15-, and 30-minute cold plasma treatment with O_2_ (Fig. 2B).

**Figure 2.**
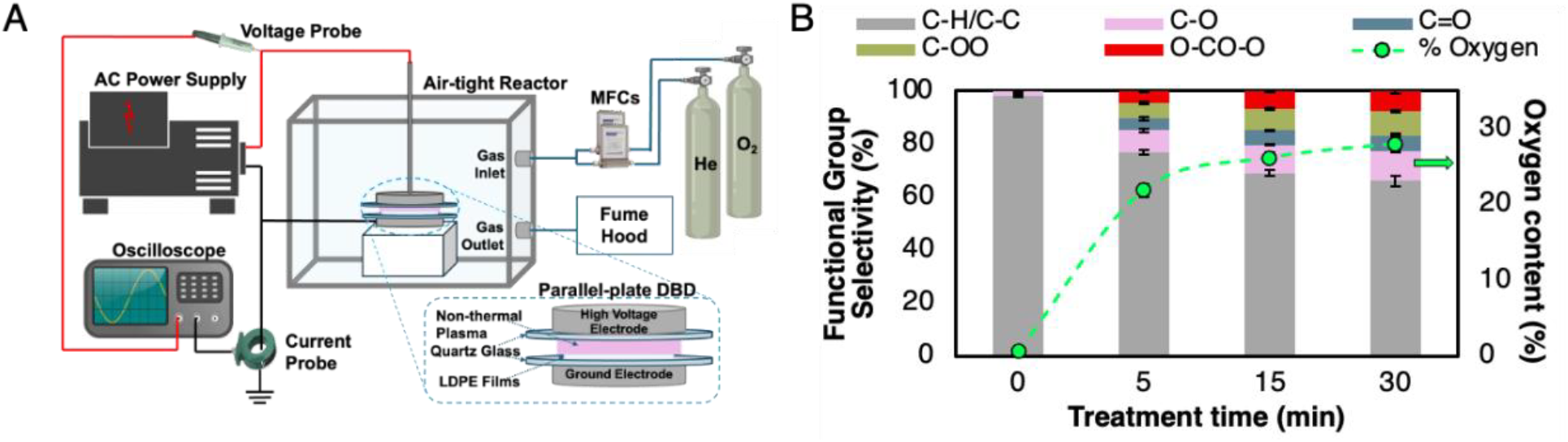
Plasma treatment was used to incorporate carbonyls onto LDPE films. (A) Schematic of entire DBD plasma reactor utilized to treat LDPE films. Red, black, and blue lines represent high-voltage electrical connections, grounded electrical connections, and gas lines, respectively. (B) XPS data of additive-free LDPE films treated with He/O_2_ plasma for 0, 5, 15, and 30 min. Bars represent chemical functional group selectivity (primary y-axis) after plasma treatment, and circles represent the percent of total oxygen content on plasma-treated LDPE films (secondary y-axis). Error bars represent the standard deviation of independent triplicate measurements.

### 3.3 Optimization of NBD-hydrazine assay on plasma-oxidized LDPE films leads to a reliable high throughput screening assay

Assay optimization was carried out to determine appropriate conditions for maximizing aldehyde detection signal on plasma-oxidized LDPE films, while limiting signal variance. NBD-H reactions with plasma-oxidized LDPE films reached completion after 40 min, evidenced by a signal plateau between 40 and 60 min of reaction time (Fig. 3A). This longer reaction time of insoluble LDPE (40 min) relative to soluble substrates (5 min)[22] is likely due to mass transfer limitations at the interface [24]. LDPE film washing was also required post-reaction to eliminate autofluorescence of residual acetonitrile/NBD-H mixture on the film and walls of the 96-well plate. Two washes with acetonitrile, performed simply by discarding reaction fluid and introducing new acetonitrile to each well with light pipette mixing, were sufficient to eliminate residual autofluorescence and signal noise (Fig. 3B). Lastly, we found that LDPE films must be oriented correctly to give appropriate signal on oxidized LDPE. When films are cut into shapes larger than the well in which the reaction will take place, assay noise greatly increases (Fig. 3C, Supplementary Fiig. 2), likely due to scattering of the fluorescent signal as the film surface is not flat and perpendicular to the detector. Therefore, LDPE films were cut into 7 mm diameter circles that are slightly smaller than the size of assay plate wells to limit assay noise (Fig. 3C).

**Figure 3.**
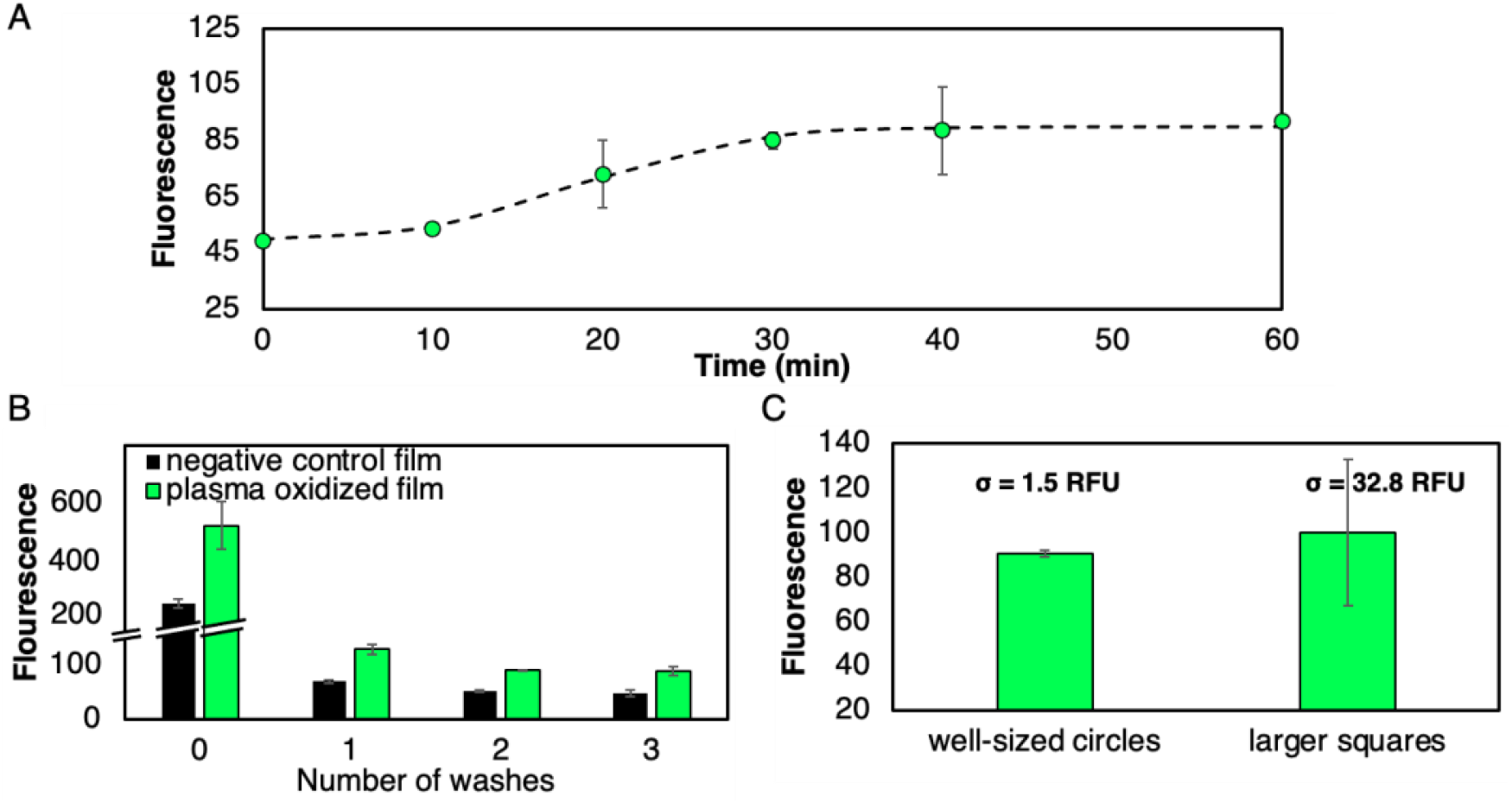
NBD-H LDPE assay optimization. (B) Kinetic NBD-H experiment on plasma oxidized LDPE films. Dashed line represents a best fit sigmoid function of the data. (C) Effect of acetonitrile washing on oxidized LDPE films post NBD-H reaction. (D) NBD-H hydrazine signal after reaction with LDPE films using 7 mm diameter circular films or 6 mm × 6 mm square films. Error bars represent the standard deviation (σ) of independent triplicate measurements.

This protocol was used on positive control plasma-oxidized LDPE films to confirm that aldehydes can be detected on the LDPE surface. Carbonyl content on plasma-oxidized LDPE was confirmed via FTIR and demonstrated that carbonyl index increases as a function of plasma oxidation treatment time (Fig. 4A). NBD-H fluorescent signal increases after plasma treatment (Fig. 4A). and this signal correlates well (R^2^ = 0.97) with carbonyl index, indicating that NBD-H can accurately and quantifiably detect carbonyls on plasma-oxidized LDPE films (Fig. 4B).

**Figure 4.**
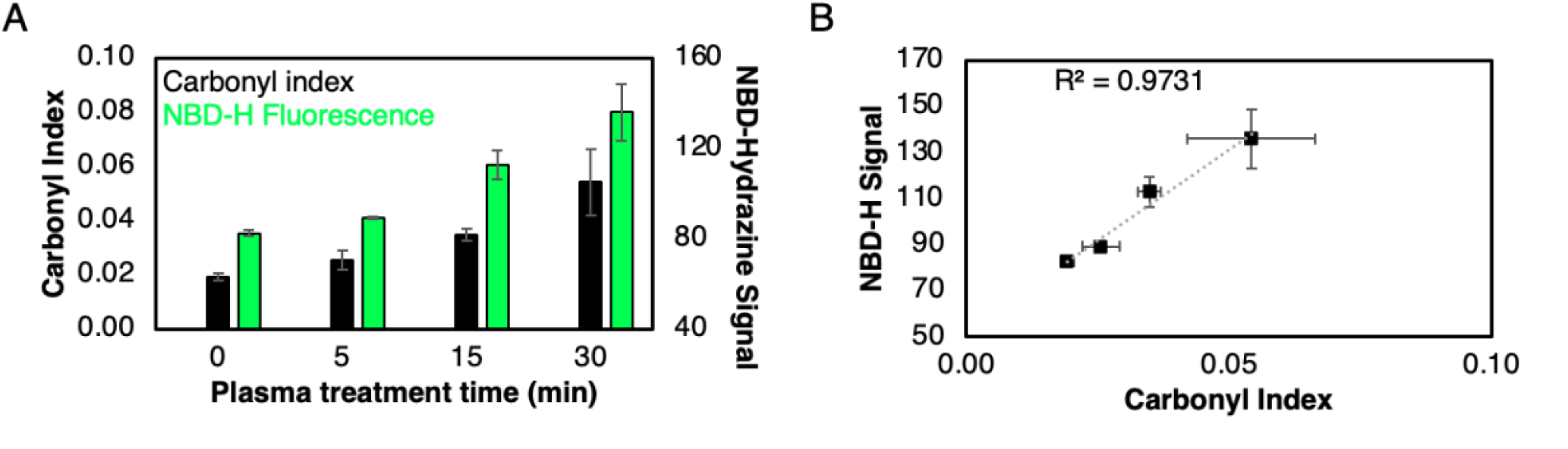
NBD-H LDPE assay optimization. (A) Carbonyl content measured via carbonyl index from FTIR spectra and NBD-hydrazine signal on stripped LDPE films. Carbonyl indices (black bars, primary y-axis)) were measured after 0-, 5-, 15-, and 30-min plasma oxidation treatment on triplicate films and NBD-H signal (green bars, secondary y-axis) was measured triplicate films after 40 min of reaction with NBD-hydrazine. (D) Carbonyl index and NBD-H signal demonstrate a strong correlation of R^2^ = 0.9731. Error bars represent the standard error of independent triplicate measurements and R^2^ values were calculated via linear regression.

### 3.4 NBD-hydrazine reliably detects enzymatic LDPE oxidation by DyP peroxidases

To evaluate the ability of our NBD-H screen to detect aldehyde formation induced via enzymatic oxidation, we tested films stripped of additives with previously characterized LDPE oxidizing DyPs that generate LDPE aldehydes as a major product.[3] Enzymes were dosed onto LDPE films in a 96-well plate and all liquid was allowed to evaporate to ensure complete surface contact of the enzyme on the LDPE film. After incubation, overnight films were rinsed via pipette mixing with 70% ethanol to remove all contamination biomolecules that could interfere with the NBD-H signal. Active enzyme preparations showed a statistically significant 140% increase relative to no-enzyme and inactivated enzyme controls (p < 0.05; one-tailed, heteroscedastic t-test; Fig. 5A, Supplementary Fiig. 3). Moreover, two doses of DyP were sufficient to bring the reaction to completion, evidenced by no change in signal after a third dose (Fig. 5B). Though only two doses are required for DyP, the number of required doses may differ between enzymes due to differences in processivity and substrate binding efficiency.

**Figure 5.**
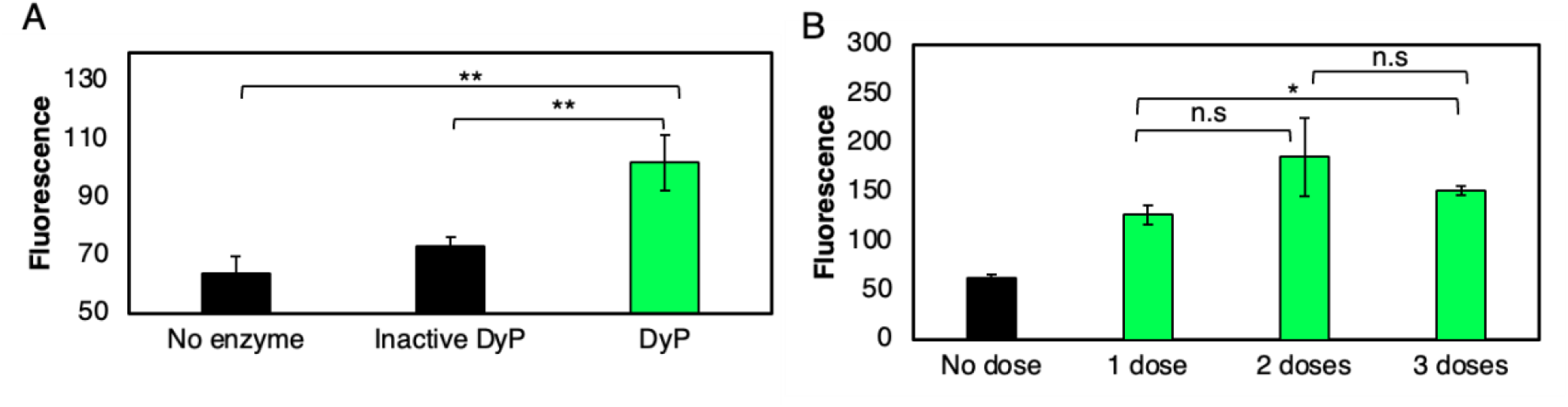
NBD-H detects aldehydes on enzymatically oxidized films. (A) NBD-H assay performed on LDPE films treated with no enzyme, β-mercaptoethanol inactivated DyP, and DyP. (B) NBD-H signal as a function of enzyme dosing, where a fresh dose was added to each film after complete drying, approximately 8 hours in assay plates. Error bars represent the standard error of independent triplicate measurements. Statistical tests were performed using a one-tailed, heteroscedastic t-test. ** represents a p-value < 0.05, * represents a p-value < 0.10, and n.s represents a p value > 0.10.

Once we validated the use of DyP on the NBD-H screening assay, we demonstrated that the assay can be used in high throughput, testing 7 DyPs homologs and inactivated controls in biological triplicate (45 total samples) in parallel on a microtiter plate with commercially available additive-free LDPE films, which we use to prevent assay interference and the generation of false positives. The high-throughput NBD-H LDPE screening platform reliably detected aldehydes on enzymatically oxidized LDPE films (Fig. 6A). This activity detection is evidenced by the statistically significant (p < 10^-6^) difference between signal from active enzymes (89 ± 32 RFU) and from denatured DyP peroxidases or inactive DyP variants (38 ± 6 RFU). Aldehyde formation was initially characterized via FTIR and correlated with NBD-H signal generation. These two independent measures were correlated (R^2^ = 0.7859) confirming that our NBD-H assay could be used as a high throughput quantitative metric for activity even on solid substrates (Fig. 6B). We encourage the use of the NBD-H assay as a high throughput screening tool to narrow the pool of enzymes to be tested by evaluating positive signals on up to thousands of enzymes tested in parallel in microtiter plates. Top performers from the NBD-H assay should then be analyzed using more rigorous oxidation detection methodologies such as FTIR, x-ray photoelectron spectroscopy (XPS), or nuclear magnetic resonance (NMR) spectroscopy to detect non-aldehyde carbonyl groups such as ketones, carboxylic acids, and esters formed as a result of enzymatic PE oxidation[16].

**Figure 6.**
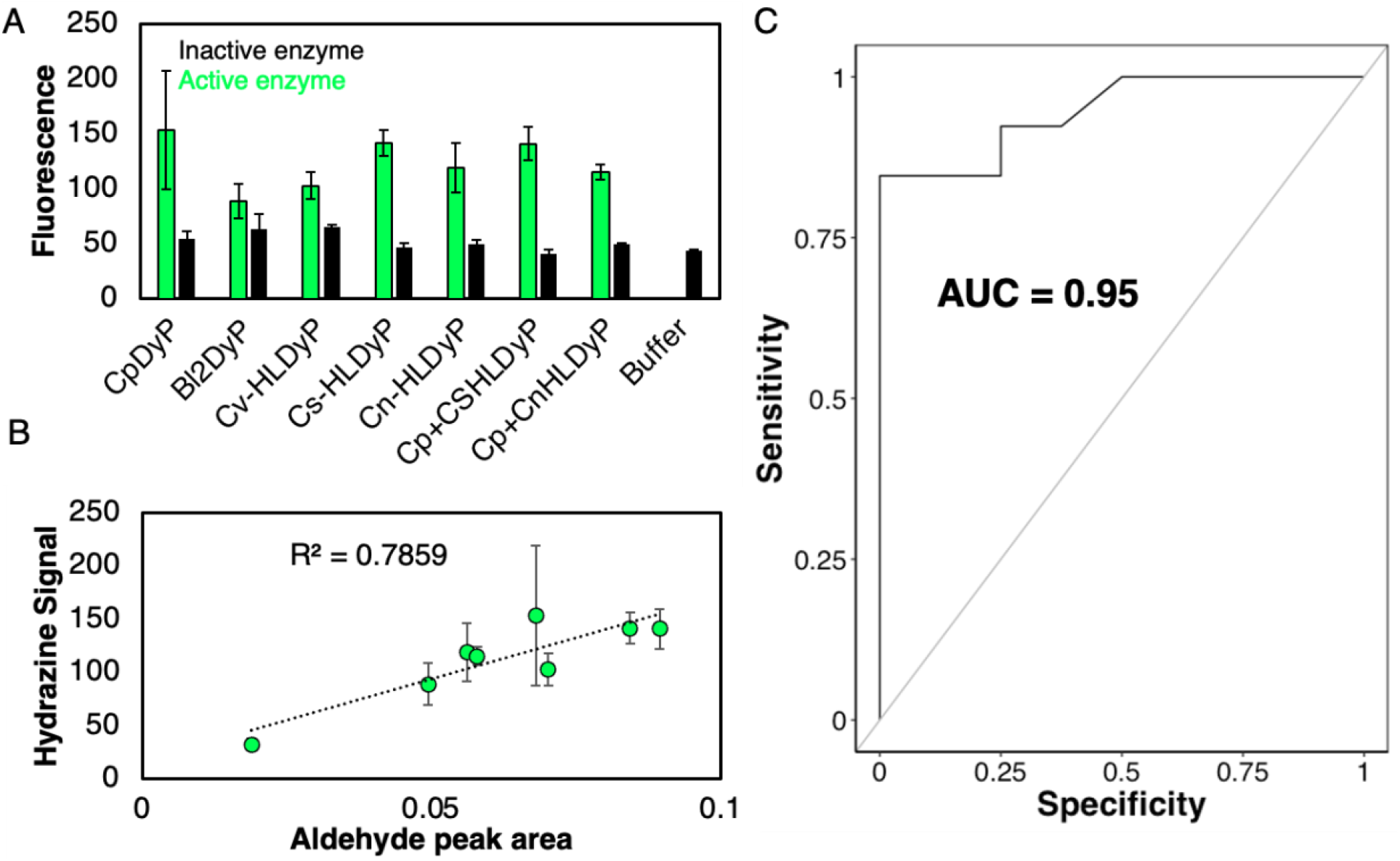
NBD-H can be used to reliably detect aldehydes on enzymatically oxidized films. (A) NBD-H fluorescent signal from an assay performed on a handful of DyP enzymes with varying activity. (B) Linear regression showing correlation between NBD-H signal and aldehyde content in DyP-treated LDPE films. Aldehyde peak area was calculated by taking the area under the aldehyde peak from FTIR spectra (centered at 1737 cm^-1^). Error bars represent the standard error of biological triplicate measurements. Aldehyde peak area and NBD-H signal in (B) represent the average of triplicate biological replicates. (C) ROC curve evaluating the performance on 50% of the data in (A). The other 50% of the data were used to train the logistic regression model to develop probabilities for fluorescence values ranging from 0 to 300, with a threshold of 52 for active vs. inactive enzyme. The area under the curve was calculated to be 0.95.

We also evaluated the ability of the assay NBD-H screening assay to be used as a simple binary classifier of LDPE active enzymes by generating a receiver operating characteristic (ROC) curve and calculating the area under the curve (AUC)[28]. The NBD-H data in Figure 6A yielded an AUC of 0.95 (Fig. 6C), meaning that the assay is considered an outstanding diagnostic test for enzyme activity[28]. Half of all datapoints (excluding the buffer only samples) in Figure 6A were used as a training set, to determine relative probabilities of RFU values in the range of ‘possible’ values from the minimum recorded value of 27 RFU to the maximum recorded value of 158 RFU. The remaining half of the dataset was used as the test dataset, with a threshold of 52 RFU delineating positive hits and negative hits as the optimum delineation point by the ROC script. ROC/AUC analysis confirms that the assay can be used to reliably detect if an enzyme is active by confirming a low rate of false positive scores greater than 52 RFU from inactive enzymes.

## 4. Discussion and Conclusions

Very few enzymes that are active on LDPE have been identified[2] due to reliance on time-consuming material characterization techniques such as FTIR that are serial in nature. Therefore, there is a need for a high throughput screening method to more quickly evaluate candidate LDPE-active enzymes to accelerate their discovery rate. Here, we directly address this need by showing that NBD-H can be used to measure aldehyde formation from enzymatic LDPE oxidation, the first step of proposed polyolefin deconstruction pathways[16–18,29]. We validate the LDPE oxidation detection capability of the developed NBD-H assay by confirming positive signal on LDPE films treated with LDPE-active DyPs[3] relative to inactive DyPs. This microtiter plate assay allows for testing of hundreds to thousands of enzymes in parallel and can be performed with an automated liquid handler to automate the enzyme discovery process. Thousands of enzymes can be screened within 24 hours through the use of this microtiter-based assay, relative to the tens of enzymes that can be tested in the same amount of time on FTIR by taking each individual spectrum in series. Moreover, the automatable nature of the assay means that individual researchers are only required to spend time setting up the liquid handling system, and the remainder of the ∼24 hours to collect thousands of data points are entirely autonomous, contrary to traditional spectroscopic methods like FTIR that require constant user interaction.

The high-throughput screening assay was developed using NBD-H due to its high selectivity for aldehydes[24]. Analogous chemical probes that react with carbonyls of all classes such as 2,4-dinitrophenylhydazine (DNPH) are not selective enough, reacting with carbonyl groups on proteins or other biomolecules in solution[30]. We show that NBD-H is appropriate for enzyme reactions with purified proteins and does not react with residual protein, as inactivated enzyme samples yield very little to no signal change relative to untreated films. Therefore, we encourage the use of the NBD-H assay as a preliminary screening tool to delineate whether or not an enzyme has oxidative activity on a polyolefin. Positive hits from the NBD-H assay should be screened using more robust oxidative metrics such as FTIR XPS, and NMR to confirm the extent of polyolefin oxidation activity by quantifying hydroxyl, ketone, and ester formation[3,10].

Since the NBD-H assay is selective for aldehydes, additional fluorometric probes can be developed for the detection of other carbonyl or alcohol groups that form as a result of enzymatic polyolefin oxidation. For example, para-methoxy-2-amino benzamidoxime has been synthesized and demonstrated to be selective for ketones[31]. Such a compound could be used in parallel with NBD-H to probe for ketone formation as well as aldehyde formation. Depending on downstream use cases for oxidized plastics, the assays could be used in parallel to select for enzymes that form more aldehydes or more ketones on the polyolefins. NBD-H and analogous probes can be used to detect oxidation of any non-hydrolysable plastic, given the similar backbone chemistries that lack carbonyl groups. Additionally, a fluorometric readout, as opposed to colorimetric, allows for use in cell sorting-based microbial assays where microorganisms from microbial communities can be sorted and subsequently screened for polyolefin oxidation in microdroplets and sorted via hydrazine signal[32]. The development of high-throughput assays such as the NBD-H assay can help greatly reduce the number of enzymes or microorganisms to be tested from large omics datasets or existing proteins in the natural sequence space. By generating such high throughput screening tools, researchers can screen exponentially more enzymes with improved reliability relative to existing indirect measurements, leading to rapid progress to discover enzymes capable of deconstructing each plastic type, paving the way for enzymatic plastic deconstruction to be feasible at scale.

## Acknowledgements

This research is supported by the U.S. Department of Energy, Office of Science, Office of Biological and Environmental Research under Award Numbers DE-SC0022018 and DE-SC0023085. This research is supported as part of the Center for Plastics Innovation, an Energy Frontier Research Center funded by the U.S. Department of Energy (DOE), Office of Science, Basic Energy Sciences (BES), under award DE-SC0021166 (HT-SEC and substrate production). RRK was funded in part by the Delaware Environmental Institute, University of Delaware and the Chemistry Biology Interface at the University of Delaware, under NIH training grant T32GM133395. The plasma work of DKN was supported by the U.S. National Science Foundation under award number 2134471. Reagents for this research were ordered in part with funds from a QIAGEN Young Scientist Research Grant award to RRK.

We thank the Advanced Materials Characterization lab at the University of Delaware for FTIR instrument time. XPS analysis was performed with the instrument sponsored by the National Science Foundation under grant No. CHE-1428149.

## Author Contributions

Ross R. Klauer: co-led all experimental work, investigation, methodology, validation, formal analysis, writing – original draft

Mekhi Williams: co-led all experimental work, investigation, methodology, validation, formal analysis, writing – original draft

Darien K. Nguyen: Investigation, writing – original draft

Megan Tarr: investigation

Dionisios G. Vlachos: reviewed and edited the manuscript

Kevin Solomon and Mark Blenner: conceptualized the study, supervised the study, acquired funding, and reviewed & edited the manuscript.

