## Supplementary information file for "A high throughput assay to detect enzymatic polyethylene oxidation"

*Supplementary Information for*: A high throughput assay to detect enzymatic polyethylene oxidation

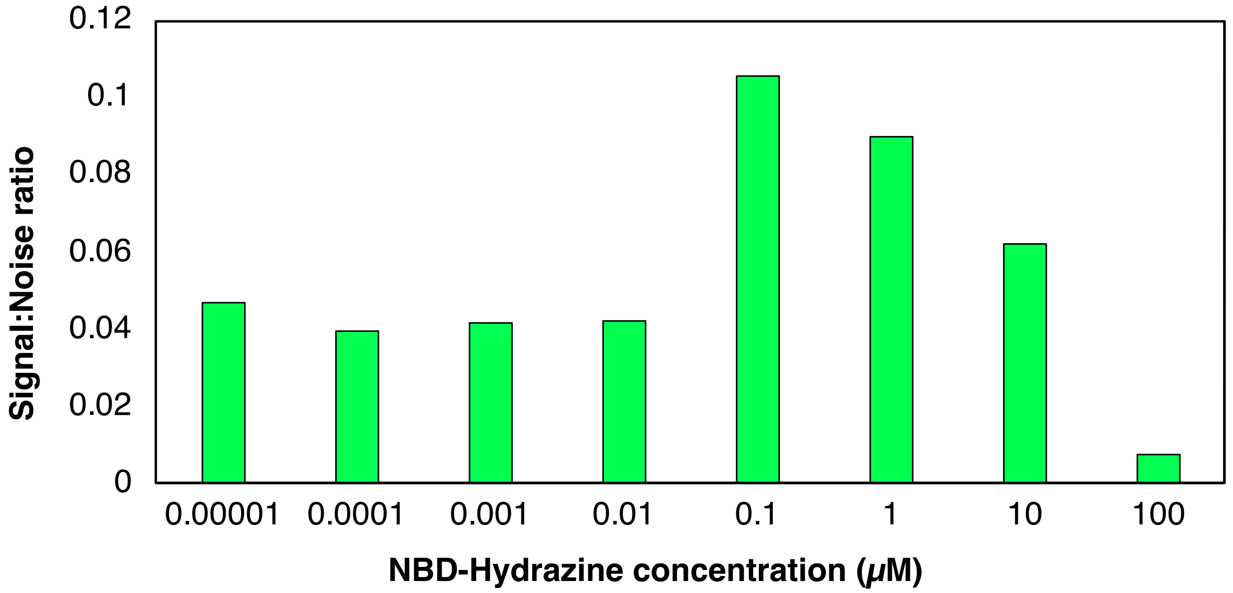


**Supplementary Figure 1: NBD-Hydrazine concentration optimization signal: noise ratio.** Signal to noise ratio was calculated using equation S1 below on the data shown in Figure 3A, to minimize false positive signal from 3-pentanone while maximizing fluorescent hydrazone signal from NBD-H reaction with pentanal.

Equation S1:

$$\frac{\frac{Pentanone_{signal}}{Pentanal_{signal}}}{baseline_{signal}}$$

Where $baseline_{signal} = average(Pentanone_{45ppm}, Pentanone_{4.5ppm})$

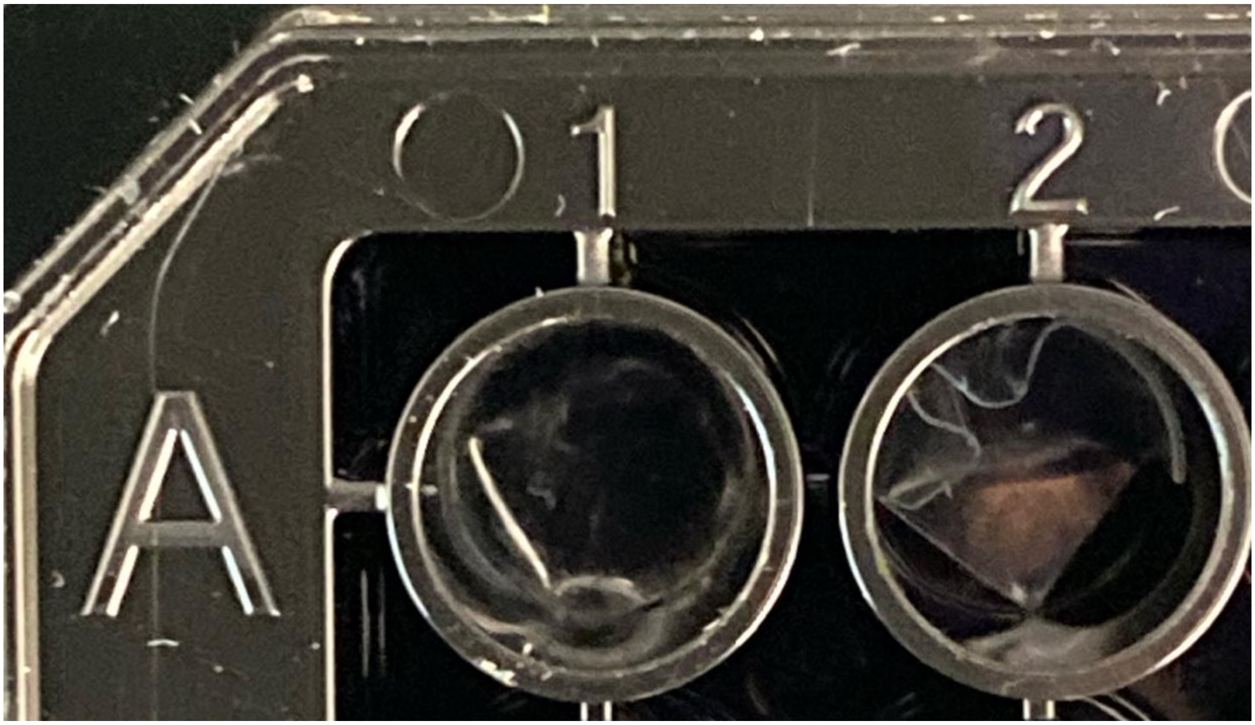


**Supplementary Figure 2: Overview of correct and incorrect film orientation for NBD-H LDPE assays.** The film on the left, in well A1 was correctly cut, using the bore end of a 200 µL pipette tip. The film on the right, in well A2 was cut as a 5 mm x 5 mm square that leads to high error in the NBD-H assay.

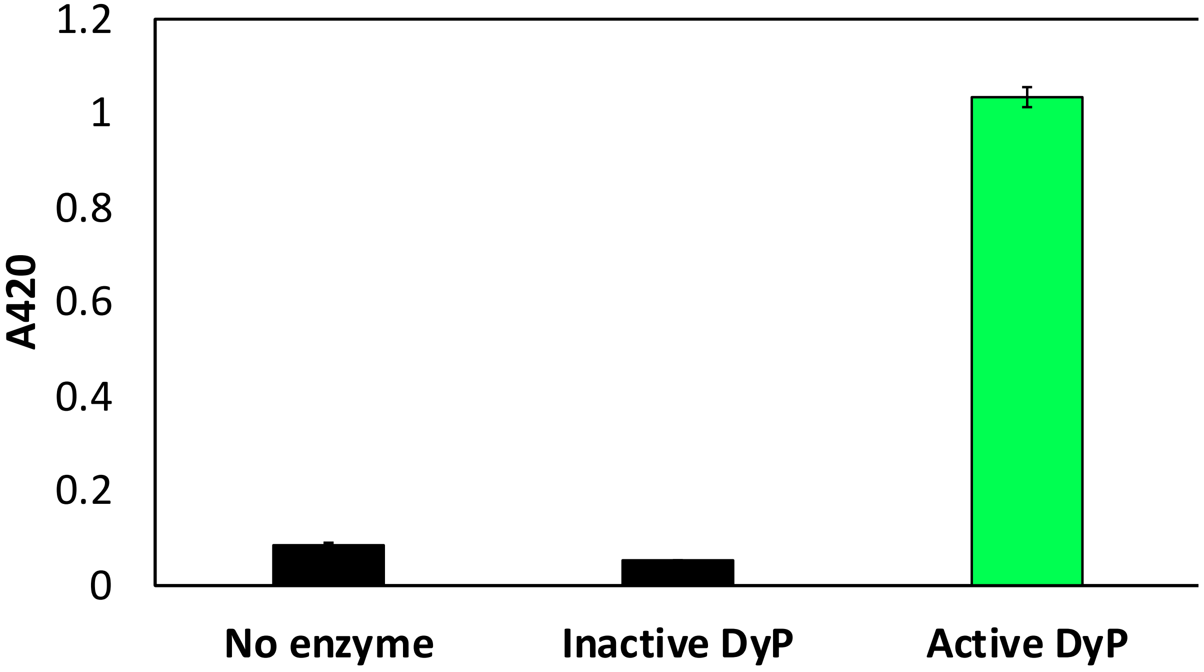


**Supplementary Figure 3: Pyrogallol enzyme activity confirming DyP inactivation by incubation with b-mercaptoethanol.** Activity of CsDyP on model substrate pyrogallol, as previously described [1], where inactive DyP was heated to 95ºC for 15 minutes in the presence of 0.35 M b-mercaptoethanol.

[1] R.R. Klauer, D.A. Hansen, Z.O.G. Schyns, L.O. Monteiro, J.A. Moore-Ott, M. Williams, M. Tarr, J. Singh, A. Mhadeshwar, L.T.J. Korley, K.V. Solomon, M.A. Blenner, Biological polyethylene deconstruction initiated by oxidation from DyP peroxidases, (2025) 2025.02.27.640435. https://doi.org/10.1101/2025.02.27.640435.
